# Reducing shock imminence eliminates poor avoidance in rats

**DOI:** 10.1101/2020.01.30.927152

**Authors:** Lindsay C. Laughlin, Danielle M. Moloney, Shanna B. Samels, Robert M. Sears, Christopher K. Cain

**Affiliations:** NYU School of Medicine, Department of Child & Adolescent Psychiatry, New York, NY 10016; Nathan Kline Institute for Psychiatric Research, Emotional Brain Institute, Orangeburg, NY 10962

## Abstract

In signaled active avoidance (SigAA), rats learn to suppress Pavlovian freezing and emit actions to escape threats and prevent footshocks. SigAA is critical for understanding aversively-motivated instrumental behavior and anxiety-related active coping. However, with standard protocols ∼25% of rats exhibit high freezing and poor avoidance. This has dampened enthusiasm for the paradigm and stalled progress. We demonstrate that reducing shock imminence with long-duration warning signals leads to greater freezing suppression and perfect avoidance in all subjects. This suggests that instrumental SigAA mechanisms evolved to cope with distant harm and protocols that promote inflexible Pavlovian reactions are poorly-designed to study avoidance.

---

In the signaled active avoidance paradigm (SigAA), rats learn to suppress Pavlovian reactions (e.g. freezing) and emit instrumental actions (e.g. shuttling) to escape warning signals (WSs) and prevent painful unconditioned stimuli (USs, typically footshocks). Understanding the psychological and neural mechanisms of SigAA is critical for several reasons. It is the prototypical paradigm for studying aversively-motivated instrumental actions (Cain et al., 2013; Rescorla and Solomon, 1967). Maladaptive or excessive avoidance responses (ARs) contribute to every major anxiety disorder (APA, 2013). Lastly, adaptive ARs reduce emotional reactions and give subjects control over environmental threats (Boeke et al., 2017; Cain and LeDoux, 2007; Choi et al., 2010; Kamin et al., 1963), suggesting a potential role in proactive coping behaviors and resilience in humans (Collins et al., 2014; LeDoux and Gorman, 2001; Moscarello and Hartley, 2017; van der Kolk, 2006).

Despite its importance as a fundamental learning mechanism with clinical relevance, SigAA research has lagged behind research on Pavlovian threats and appetitive instrumental behavior (Cain, 2019; Krypotos et al., 2015; LeDoux et al., 2017). The phenomenon of “poor avoidance” has been one major obstacle to progress. Avoidance acquisition is typically slower than Pavlovian conditioning, but most animals learn to prevent >80% of scheduled shocks in two-way shuttlebox tasks. However, a significant subset of animals exhibit high freezing and rarely emit ARs (Choi et al., 2010; Galatzer-Levy et al., 2014; Keehn, 1967; Martinez et al., 2013). For some tasks, poor avoidance is the rule rather than the exception (Neffinger and Gibbon, 1975; Solomon and Brush, 1954). From a practical standpoint, avoidance studies are more costly and time-consuming because poor avoiders must be replaced. Pretraining loss-of-function studies are also ill-advised with SigAA, since there is no reliable way to predict which animals will acquire ARs. Finally, the poor avoidance phenomenon raises questions about whether instrumental AR learning is a major component of defense worthy of study (Bolles, 1975; Fanselow, 1997; 2018). Animals evolved defensive mechanisms to cope with predators, not shocks, and it is difficult to see how a trial-and-error learning mechanism that often fails could have evolved under predatory pressure.

One simple explanation is that researchers have used suboptimal protocols for studying avoidance in the laboratory. SigAA is typically evaluated in small chambers with short-duration WSs and high-density shock protocols. These conditions are ideal for modelling a state of high predatory imminence that triggers hard-wired, stereotyped fear-like reactions (e.g. freezing) that are incompatible with ARs (Fanselow and Lester, 1988; Weiss et al., 1968). However, prey animals spend much more time in a state of low predatory imminence where encounters with predators are temporally distant or uncertain. Perhaps instrumental avoidance mechanisms evolved to deal with these anxiety-like states, where animals must balance conflicting needs (Cain, 2019; Diehl et al., 2019; Gray and McNaughton, 2000). Under these “pre-encounter” conditions, less rigid defensive behaviors may not interfere with AR learning.

To solve the poor avoidance problem and optimize avoidance training, we designed two experiments to evaluate AR learning while systematically varying threat intensity. In the first, WS-US contingency was varied to test how US certainty affects AR learning. In the second, WS duration was varied to test how US imminence affects AR learning. In Pavlovian studies, reducing US certainty or imminence appears to promote anxiety over fear; freezing reactions are diminished and more flexible antipredator strategies increase (Blanchard et al., 1989; Cain et al., 2005; Goode et al., 2019; Helmstetter and Fanselow, 1993; Kim and Jung, 2018; Mobbs et al., 2007; Rescorla, 1968; Waddell et al., 2006). Lesions that impair freezing rescue ARs in poor avoiders, suggesting that freezing reactions interfere with avoidance (Choi et al., 2010; Lazaro-Munoz et al., 2010; Moscarello and LeDoux, 2013). Pavlovian reactions also impair avoidance performance in humans (Rigoli et al., 2012). Thus, we predicted that both methods of reducing threat intensity would decrease Pavlovian freezing and improve AR learning.

Experiments were conducted on adult male and female Sprague-Dawley rats (Hilltop Lab Animals) weighing 300-350g on arrival. Rats were pair-housed by sex, had ad lib access to food and water, and were tested during the light phase of a 12:12-hour light:dark schedule. All procedures were approved by the NKI-IACUC.

All rats received 10 days of two-way SigAA training in standard shuttleboxes equipped with speakers, houselights, cameras, grid floors and infrared beams to detect shuttling (Coulbourn Instruments). Sessions included a 5-minute acclimation followed by 15 trials where warning stimuli (80dB white noise) preceded scrambled 0.5s footshocks (males: 1.0mA, females: 0.7mA). Session 1 always began with an inescapable Pavlovian trial, ensuring that all subsequent WS-shuttles occurred during threat of shock. For all subsequent trials, shuttling to the opposite chamber side terminated the WS, produced feedback (5s, 5kHz, 80dB tone), and cancelled the upcoming shock (if scheduled). Shuttling was automatically recorded by Graphic State software (Coulbourn Instruments) and freezing was recorded to video files for off-line analysis. Intertrial intervals (ITIs) averaged 2-minutes unless otherwise stated. Avoidance percentage was calculated for individuals each session [(WS-shuttles/Trials)*100]. Avoidance latency reflects the time from WS onset to shuttle, with failures recorded as the full WS duration. Freezing was scored during the WS for select sessions by two experienced raters blind to group (inter-rater reliability correlation >0.9). To facilitate comparisons of freezing suppression between studies, Session 10 freezing was also analyzed as a percentage of Session 1 freezing (calculated for individuals and then averaged).

In Experiment 1, rats received SigAA training with a standard 15s WS except the likelihood of receiving a shock on failure trials was varied (100%, 50% or 25%; Figures 1A-B). Two-way ANOVAs (GraphPad Prism v8) indicate group differences in acquisition rate for both AR% and AR latency (Group x Session interactions: F_(18,189)_=1.8, p=0.02; F_(18,189)_=2.0, p=0.01), however, reducing WS-US contingency did not improve learning. These differences appear to be driven by a deficit in the 25% group, where rats shuttled on average more slowly and less frequently. On average, freezing declined across avoidance training but there were no significant group differences. A two-way ANOVA revealed a significant effect for Session (F_(2,42)_=21.6, p<0.01), but not for Group (F_(2,21)_=0.26) or the Group x Session interaction (F_(4,42)_=1.7; Figure 1C). Similarly, Session 10 freezing was lower on average than Session 1 but no group differences were observed (Figure 1D; One-way ANOVA: F_(2,21)_=2.1).

**Figure 1.**
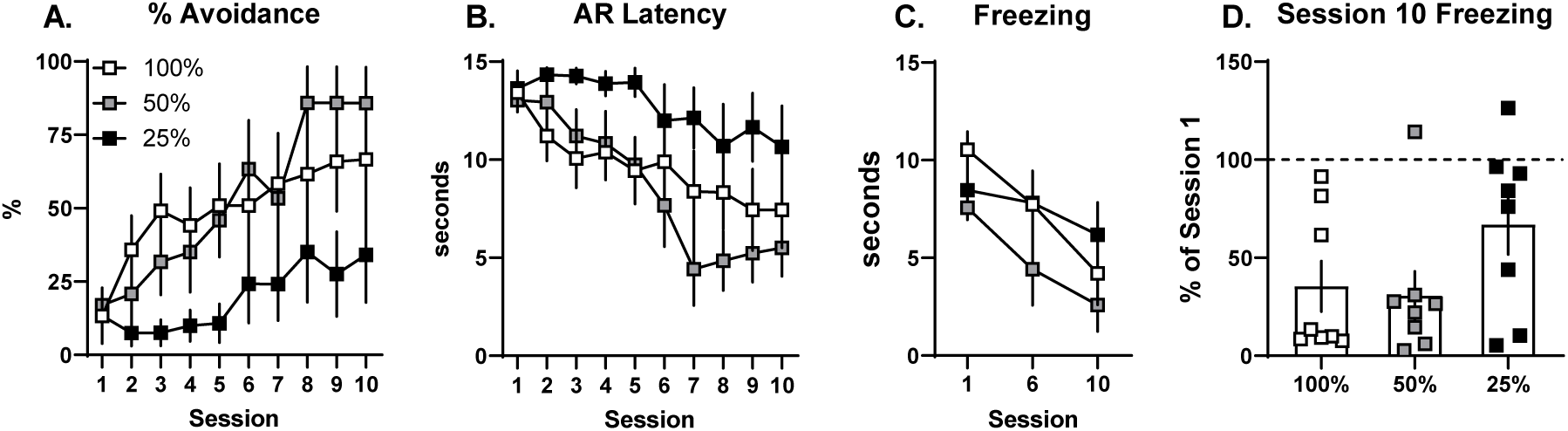
Reducing WS-US contingency does not improve avoidance. **A)** Mean percent avoidance by session. **B)** Mean avoidance response (AR) latency by session. **C)** Mean seconds freezing during warning signals for Sessions 1, 6 and 10. **D)** Mean Session 10 freezing expressed as a percentage of Session 1 freezing. Squares represent individuals. N=8/group (4 females, 4 males). Error bars = S.E.M. *p<0.05 vs. 100% WS-US contingency group

In Experiment 2, rats received avoidance training with 100% WS-US contingency (shock delivered on every failure trial), but the WS duration was varied (15s, 60s or 240s; Figures 2A-B). AR% increased across training (Session: F_(9,189)_=21.1, p<0.01) and there was a significant effect of WS duration (Group: F_(2,21)_=4.7, p=0.02), however the pattern of change over time did not differ between groups (Group x Session: F_(18,189)_=0.9). The effect of WS duration was driven mainly by the 240s group, where AR% was higher than the 15s group for Sessions 2-5 (Dunnett’s tests). Remarkably, every rat in the 240s group showed perfect avoidance from Session 3 until the end of training (no failures). As expected for different WS durations, large differences in AR latency were observed across training (Group x Session: F_(18,189)_=21.9, p<0.01). These differences are not very informative early in training when failures were common and WS-duration determined AR latency. Interestingly, AR latencies were very similar by the end of training, even though rats in the 60s and 240s groups had much more time to emit ARs (Figure 2B, inset). Large freezing differences were also apparent across training (Figure 3C; Groups x Session: F_(4,42)_=37.7, p<0.01). This may be partly explained by the different WS durations. Rats in all groups froze for most of the WS early in training and freezing declined to similarly low levels as ARs were acquired. Dunnett’s post tests revealed that rats in the 240s and 60s groups froze more than rats in the 15s group during Session 1 only (p values <0.01). Rats in the 240s group showed the strongest suppression of freezing by Session 10 (Figure 2D; one-way ANOVA: F_(2,21)_=3.4, p=0.05; Dunnett’s test vs. 15s group: p<0.05). This appears to reflect more than the programmed differences in WS duration; unlike the other groups, no rats in the 240s group maintained or increased freezing across training (as occurs in poor avoiders; Lazaro-Munoz et al., 2010).

**Figure 2.**
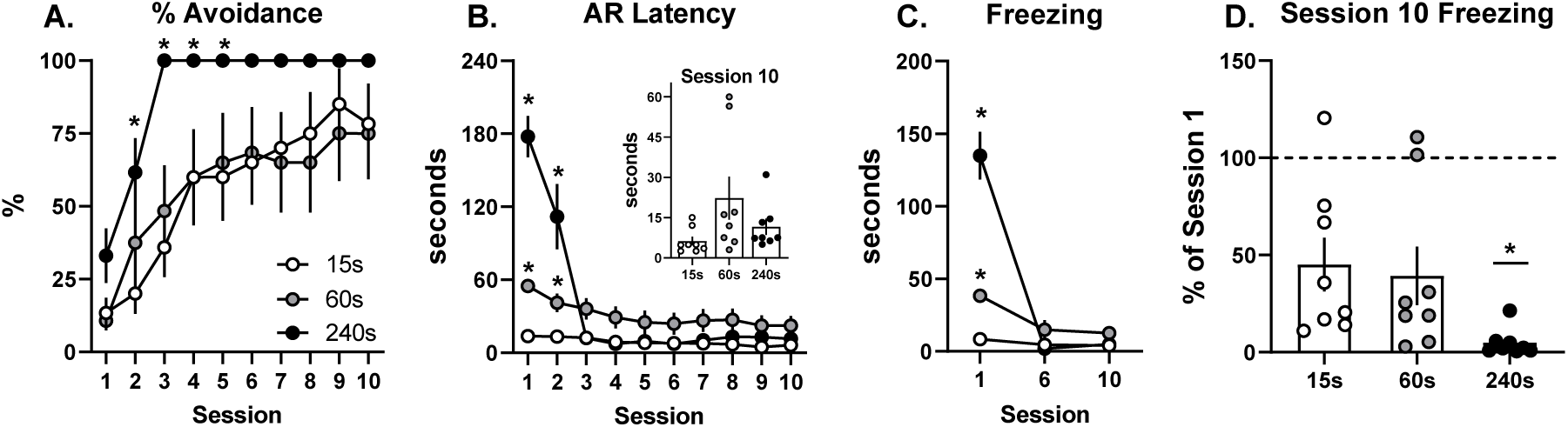
Reducing US imminence leads to perfect avoidance. **A)** Mean percent avoidance by session. **B)** Mean avoidance response (AR) latency by session. ***inset*:** mean AR latency for individuals during Session 10. **C)** Mean seconds freezing during warning signals for sessions 1, 6 and 10. **D)** Mean Session 10 freezing expressed as a percentage of Session 1 freezing. Dots represent individuals. N=8/group (4 females, 4 males). Error bars = S.E.M. *p<0.05 vs. 15s-WS group

**Figure 3.**
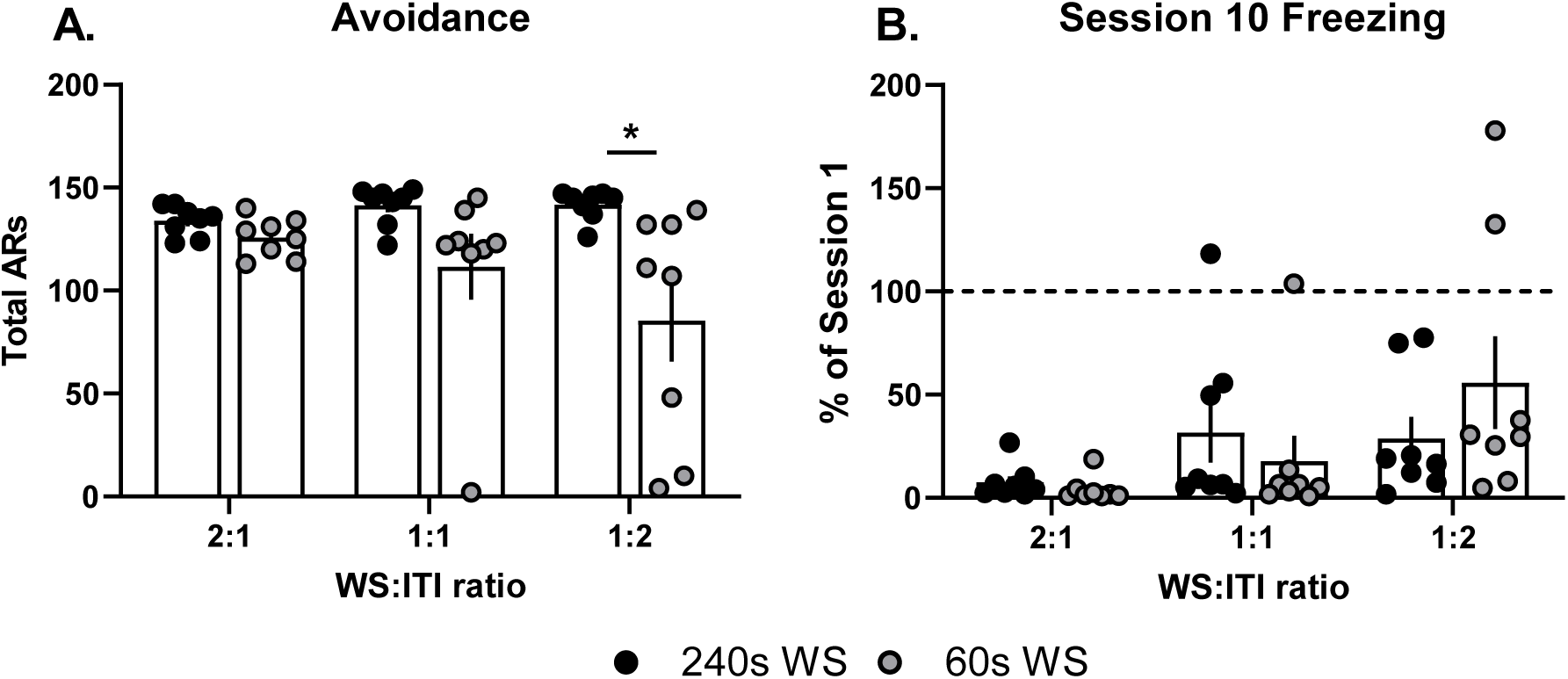
Reducing the WS:ITI ratio fails to impair avoidance with a 240s warning signal. A) Total avoidance responses emitted across 10 sessions of training. B) Mean Session 10 freezing expressed as a percentage of Session 1 freezing. N=8/group (4 females, 4 males). Bars represent separate groups. Bar height indicates group mean. Dots represent individuals. Error bars = S.E.M. *p<0.05 for 240s vs. 60s WS groups

One potential criticism of the long WS is that apparent ARs reflect locomotor activity not instrumental shuttling. To address this, we replicated AR training with the 240s WS (n=8; 5 females, 3 males) and included Yoked controls (n=8; 4 females, 4 males). All rats in the 240s-WS group again attained perfect avoidance (not shown). Yoked controls shuttled far less frequently during the WSs than Master rats (<0.2 shuttles/trial on average by training end; Group x Session interaction (F_(9,126)_ = 8.3, p<0.01). This supports the notion that WS shuttles represent instrumental ARs even with long-duration WSs.

Experiment 2 was designed to test the effect of US imminence on avoidance learning. However, because WS duration was varied while the ITI was held constant, another explanation is possible. In Pavlovian studies, the conditioned stimulus (CS) to ITI ratio has a strong impact on performance of Pavlovian reactions (reviewed in Balsam et al., 2010). Specifically, higher CS:ITI ratios weaken responding, perhaps because the signal loses informational value relative to the background context (Gibbon and Balsam, 1989). Thus, it is possible that our long WS enhanced avoidance because it increased the WS:ITI ratio and weakened competing freezing reactions. To address this, we evaluated avoidance and freezing with WS:ITI ratios of 2:1, 1:1 and 1:2 using two different WS durations (60s and 240s). We trained four new groups of rats: 60s-WS:30s-ITI (2:1), 60s-WS:60s-ITI (1:1), 240s-WS:480s-ITI (1:2), and 240s-WS:240s-ITI (1:1). The remaining groups for the analysis came from Experiment 2: 60s-WS:120s-ITI (1:2) and 240s-WS:120s-ITI (2:1). Figure 3A depicts total (cumulative) ARs across training. A two-way ANOVA revealed a significant effect of WS-duration (F_(1,42)_=12.9, p<0.01) and a non-significant trend towards a WS-duration x Ratio interaction (F_(2,42)_=2.6, p=0.09). The main effect for Ratio was not significant (F_(2,42)_=1.3). Thus, reducing the WS:ITI ratio failed to impair ARs and perfect avoidance was achieved for every rat trained with the 240s-WS. The same manipulation may reduce avoidance with a shorter WS; total ARs declined as the WS:ITI ratio dropped for the 60s WS. Further, in the 2:1 condition, rats in the 60s group avoided less than rats in the 240s group (planned Sidak’s comparison). Suppression of freezing was more sensitive to the WS:ITI ratio (Figure 3B). Session 10 freezing increased as the WS:ITI ratio dropped (Ratio: F_(2,42)_=0.03, p=0.03), but this effect was not modulated by WS-duration (WS-duration: F_(1,42)_=0.09; WS-duration x Ratio: F_(2,42)_=1.4). This suggests that the WS:ITI ratio is not a major determinant of AR acquisition with very long WSs. However, there are indications that reducing this ratio promotes freezing and impairs AR learning with shorter WSs. The reverse also appears true; using a 2:1 ratio led to very low Session 10 freezing and perfect avoidance for 7 of 8 rats trained with the 60s-WS. Exploring a wider range of WS:ITI ratios may help clarify these findings.

Lastly, all experiments included both female and male subjects. Avoidance learning with the 240s WS was nearly identical between the sexes as measured by AR% (Session: F_(9,126)_=41.0, p<0.01, Sex: F_(1,14)_=2.3, Session x Sex: F_(9,126)_=1.0) and AR latency (Session: F_(9,126)_=39.3, p<0.01, Sex: F_(1,14)_=3.6, Session x Sex: F_(9,126)_=0.6). Freezing across training was also very similar (Session: F_(2,28)_=81.5, p<0.01, Sex: F_(1,14)_=0.29, Session x Sex: F_(2,28)_=0.08). Sex differences were difficult to evaluate in the other conditions due to poor avoiders, which appear to occur randomly (equally likely in both sexes).

Our major finding is that reducing US imminence by extending WS duration greatly facilitates SigAA learning. In four separate groups trained with the long-duration WS, every rat learned and performed the task perfectly (no subsequent failures), sometimes in fewer than 30 trials. The benefits of the long-duration WS also resisted manipulations of the WS:ITI ratio that promote competing freezing reactions. Several observations also argue against the concern that shuttling during long-duration WSs reflects exploration rather than instrumental ARs. First, exploration was severely depressed early in training where rats froze for more than 60% of the WS. Second, once the response was acquired, ARs were emitted with short latencies (usually <15s). Third, yoked controls shuttled during the WS at a far lower rate than master rats.

What might explain the enhanced efficiency of SigAA with long-duration WSs? Though there are some reports of improved SigAA learning with slightly longer or more complex WSs (Archer et al., 1984; Coll-Andreu et al., 1993; Levis and Stampfl, 1972; Satorra-Marin et al., 2001; Solomon and Brush, 1954; Terburg et al., 2018), this has not been systematically studied. There are far more studies of US imminence using Pavlovian paradigms. These suggest that threats activate different components of the survival circuit depending on proximity to harm (modeled by CS-US delay; Davis, 1998; Goode et al., 2020; Mobbs et al., 2009; Mobbs et al., 2007; Sullivan et al., 2004; Waddell et al., 2006; Walker and Davis, 1997). Short-duration CSs recruit amygdala and periaqueductal gray to emit short-latency, inflexible, hard-wired responses that function to prevent threat escalation (e.g. freezing, a post-encounter response) or escape harm (e.g. flight, a circa-strike response). Long-duration CSs recruit bed nucleus of the stria terminalis and prefrontal cortex to flexibly reorganize behavior (e.g. thigmotaxis, altered meal-patterns), presumably to prevent threat escalation and prepare the organism to defend against distant or uncertain harm. Importantly, high US-imminence restricts behavior to species-specific defense reactions whereas low US-imminence balances defense with other behaviors like exploration and reward procurement (Blanchard and Blanchard, 1989; Bolles, 1970; Fanselow, 2018; Gray and McNaughton, 2000; Mobbs et al., 2015; Mobbs and Kim, 2015). Thus, long-duration WSs likely trigger less intense defensive strategies and allow for active responses like shuttling. This is consistent with observed patterns of freezing; though 240s-WS rats froze significantly early in training (∼62%), they had more time to emit the AR and experience the instrumental contingency than rats in the other groups. Freezing appeared to be more easily suppressed in this condition too. Thus, optimal trial-and-error SigAA mechanisms may have evolved under low threat conditions, where errors (failures to emit the AR) lead to more intense threats and not necessarily harm.

Though our hypothesis about lowering threat intensity to improve avoidance was supported by the US-imminence experiment, it was not supported by reducing WS-US contingency. Rats receiving shocks on only 25% of failure trials did not perform better than rats in the 100% contingency condition. We see two possible explanations for this. First, SigAA learning depends, at least in part, on omission of expected US presentations (Bolles et al., 1966; Cain, 2019; Hunter, 1935; Kamin, 1956). So even if 25% WS-US contingency reduces certainty and competing freezing reactions, this may have been offset by degradation of an important reinforcement signal. Second, 10 sessions of SigAA training may have been too few to observe the benefit of reduced WS-US contingency. Additional work is needed to clarify these points. We found only two other studies that varied shock delivery on failure trials (Boren and Sidman, 1957; Neffinger and Gibbon, 1975). Though these were conducted quite differently (contingency manipulated after standard training, poor avoiders eliminated, no-ITI protocols, bar-press avoidance etc.), both confirm that reducing the likelihood of shock on failure trials leads to an avoidance decrement.

Interestingly, our follow-up experiment suggests another possible way to improve SigAA efficiency: increase the WS:ITI ratio. Pavlovian studies show that increasing the CS:ITI ratio impairs Pavlovian reactions (Balsam et al., 2010; Delamater and Holland, 2008; Stein et al., 1958). This is likely a result of the CS losing informational value relative to the background context (Gibbon and Balsam, 1989), making the CS a weaker threat. We see a similar pattern in freezing suppression during SigAA training; increasing the WS:ITI ratio produced weaker Session 10 freezing and near-perfect avoidance with the shorter 60s-WS. If replicated, this protocol could ensure good avoidance in all subjects with significantly shorter session durations.

In summary, we describe two simple procedural methods to improve SigAA learning and eliminate poor avoidance: increase the WS duration and/or the WS:ITI ratio. This removes a major obstacle to SigAA research that has dampened enthusiasm for the paradigm over decades. Experiments requiring pre-training manipulations can be used with confidence if controls reliably learn and perform ARs. The explanation for enhanced avoidance with low-intensity threats also aligns with functional behavior systems theories of defensive behavior and Pavlovian studies of US-imminence (Fanselow and Lester, 1988; Mobbs and Kim, 2015; Waddell et al., 2006), but is inconsistent with two-factor “fear” theories of avoidance that assume avoidance positively correlates with threat intensity (Levis, 1989; Miller, 1948; Mowrer and Lamoreaux, 1946). This work may also help explain how strong avoidance responses may be acquired in human anxiety even when harm is not imminent.

## Acknowledgements

This work was supported by the National Institute of Mental Health of the National Institutes of Health under award numbers [R01MH114931] to C. Cain and [R21MH116242] to R. Sears. Additional funding for this project was provided by The William S. McIntyre Foundation to R. Sears. The authors thank Peter Balsam and Michael Fanselow for helpful discussions about the data.

